# Extracellular Vesicles From Metabolically Healthy Obesity Convey Distinct Molecular Signals That Initiate Endothelial Dysfunction: A Multi-Omics Study in Adults of African Ancestry

**DOI:** 10.64898/2026.04.09.717593

**Authors:** Malak Abbas, Camryn Bragg, Ahmed Gharib, Abdel G. Elkahloun, Merry Lindsey, Amadou Gaye

**Author notes:** Corresponding Author: Amadou Gaye, PhD, Department of Integrative Genomics and Epidemiology, Meharry Medical College 1005 Dr DB Todd Jr Blvd, Nashville, TN 37208.

## Abstract

**Background:** Metabolically healthy obesity (MHO) is unstable, with up to 80% of individuals progressing to metabolically abnormal obesity (MAO), yet mechanisms underlying this transition remain unclear. African Americans bear a disproportionate burden of obesity-related cardiovascular disease. Circulating extracellular vesicles (EVs) mediate inter-organ communication and may drive MAO-related vascular dysfunction.

**Methods:** Adults of African ancestry were classified as metabolically healthy lean (MHL, n=14), MHO (n=9), or MAO (n=16). Plasma-derived EVs were characterized and their microRNA cargo profiled. Human coronary artery endothelial cells were treated with EVs from each group to assess nitric oxide signaling, oxidative stress, inflammatory activation, and mitochondrial dynamics.

**Results:** MHO participants exhibited preserved insulin sensitivity and lower inflammation compared with MAO despite comparable adiposity. EVs from MHO carried a distinct microRNA signature enriched in miR-148a-5p, miR-181c-5p, and miR-1255a, linked to antioxidant and matrix regulatory pathways. MAO EVs were enriched in miR-3613-3p, miR-6842-3p, and miR-326, targeting inflammation and insulin resistance pathways. Compared with both MHL and MHO EVs, MAO EVs suppressed endothelial nitric oxide synthase phosphorylation and reduced nitric oxide bioavailability, with increased reactive oxygen species and ICAM-1 expression. MHO EVs induced an intermediate phenotype with disrupted mitochondrial morphology, supporting a graded continuum of endothelial stress.

**Conclusions:** MHO represents a biologically active intermediate state. Circulating EVs from MHO individuals convey molecular signals that impair endothelial and mitochondrial function, predisposing to vascular injury and progression toward MAO. EV-associated microRNAs are mechanistic mediators and candidate biomarkers of metabolic and vascular deterioration in obesity.

**CLINICAL PERSPECTIVE:** *What Is New?:*

- This study systematically investigated extracellular vesicles derived from metabolically healthy obese individuals to define direct vesicle effects on endothelial function using integrated omics coupled to functional outputs.
- Extracellular vesicles from metabolically healthy obesity convey a distinct molecular and biological signature that distinguishes lean and metabolically abnormal obesity.
- Metabolic health status, rather than obesity alone, drives extracellular vesicle-mediated endothelial nitric oxide signaling, oxidative stress, inflammation, and mitochondrial dynamics.

*What Are the Clinical Implications?:*

- These findings explain why some individuals with obesity exhibit preserved vascular function while others develop early endothelial dysfunction.
- Stratifying obesity by metabolic health status improves cardiovascular risk assessment beyond body mass index alone.
- Targeting extracellular vesicle signaling pathways represents a novel strategy to prevent metabolically healthy individuals from progressing to metabolically abnormal obesity.

## INTRODUCTION

Obesity affects more than 650 million adults worldwide and remains a leading driver of type 2 diabetes, hypertension, and cardiovascular disease.1,2 Within this spectrum, a subset of individuals with obesity, termed metabolically healthy obesity (MHO), maintains normal insulin sensitivity, lipid profiles, and blood pressure despite excessive adiposity.3,4 However, longitudinal studies demonstrate that MHO is rarely durable: up to 80% of individuals with MHO transition to metabolically abnormal obesity (MAO) within a decade, accompanied by systemic inflammation, insulin resistance, and vascular dysfunction.5–7 These findings suggest that MHO represents a transient, compensated state rather than a truly benign phenotype.

The mechanisms underlying this metabolic decline remain incompletely defined. Emerging evidence implicates circulating extracellular vesicles (EVs) as endocrine mediators of inter-organ communication in obesity.8,9 EVs are nanosized membrane-bound vesicles that transport proteins, lipids, and nucleic acids, including microRNAs (miRNAs), which regulate gene expression and cell signaling in recipient tissues.10,11 In obesity, adipose- and immune-derived EVs circulate at increased levels and carry miRNA cargo implicated in insulin resistance, inflammation, and endothelial dysfunction.12,13 Preclinical studies provide evidence of causality: transfer of EVs from obese donors to lean mice induces systemic insulin resistance and glucose intolerance, while delivery of obesity-associated miRNAs recapitulates key metabolic abnormalities, including hepatic steatosis and increased visceral adiposity.14–16 These vesicle-borne signals may therefore link adipose tissue stress to early vascular injury.

African Americans are disproportionately affected by obesity and its cardiometabolic complications, including hypertension, type 2 diabetes, and cardiovascular disease, yet remain underrepresented in mechanistic studies of obesity-related vascular dysfunction. The higher prevalence and earlier onset of metabolic abnormalities in this population underscore the need to define molecular pathways that govern the transition from healthy to unhealthy obesity in individuals of African ancestry. We therefore investigated EV-mediated molecular communication across the metabolic obesity continuum in this population. We hypothesized that circulating EV cargo differs across metabolically healthy lean (MHL), MHO, and MAO states, and that vesicles drive a graded pattern of endothelial stress. Specifically, we postulated that EVs from MHO individuals, while less pathogenic than those from MAO, already impair nitric oxide (NO) signaling and mitochondrial integrity in endothelial cells, marking the onset of subclinical vascular dysfunction.

## METHODS

An expanded Methods section, including detailed protocols for all assays and bioinformatic pipelines, is available in the Online Data Supplement. Please see the Major Resources Table in the Supplemental Materials.

### Study Population and Phenotype Classification

We studied a cohort of 39 adults of African ancestry stratified into 3 groups: metabolically healthy lean (MHL, n=14; BMI 18.5-24.9 kg/m²), metabolically healthy obese (MHO, n=9; BMI ≥30 kg/m²), and metabolically abnormal obese (MAO, n=16; BMI ≥30 kg/m²). Individuals with overweight (BMI 25.0-29.9 kg/m²) were excluded by design to ensure clear separation between lean and obese phenotypes. Group definitions (Table 1) were based on modified criteria incorporating inflammatory status, consistent with the Wildman classification.22 MHL participants had normal BMI without metabolic abnormalities; MHO participants had obesity with a metabolic profile comparable to MHL; and MAO participants had obesity with ≥2 metabolic abnormalities or established cardiometabolic disease. All participants were free of clinical cardiovascular disease and anti-inflammatory or weight-loss medications. The study protocol was approved by the institutional ethics board, and written informed consent was obtained.

**Table 1.** Criteria for Classification of Metabolic Phenotypes.

| Group | BMI (kg/m<sup>2</sup>) | Metabolic Syndrome Components | Additional Criteria |
| --- | --- | --- | --- |
| **MHL** | 18.5-24.9 | None: SBP/DBP ≤130/85 mmHg; TG/HDL ≤1.32 (W) or ≤1.65 (M); HDL ≥50 (W) or ≥40 (M) mg/dL; FG ≤100 mg/dL; HOMA-IR ≤5.1; CRP ≤0.3 | Normal insulin sensitivity; no hypertension, dyslipidemia, or hyperglycemia |
| **MHO** | ≥30 | Same as MHL | Same as MHL |
| **MAO** | ≥30 | ≥2 of the above metabolic syndrome components | May include T2D, hypertension, or dyslipidemia under treatment |

BMI indicates body mass index; CRP, C-reactive protein; DBP, diastolic blood pressure; FG, fasting glucose; HDL, high-density lipoprotein; HOMA-IR, homeostatic model assessment of insulin resistance; M, men; MAO, metabolically abnormal obese; MHL, metabolically healthy lean; MHO, metabolically healthy obese; SBP, systolic blood pressure; T2D, type 2 diabetes; TG, triglycerides; W, women.

### Plasma Preparation and EV Isolation

EDTA-anticoagulated plasma was prepared by sequential centrifugation and filtration through a 0.45-µm membrane following International Society for Extracellular Vesicles (ISEV) standardized protocols. EVs were isolated using the Norgen Biotek Plasma/Serum Exosome Purification Kit. Vesicle size and morphology were evaluated by transmission electron microscopy (TEM) and nanoparticle tracking analysis (NTA). Immunoblotting confirmed enrichment of canonical small EV markers (CD9, CD63, CD81, ALIX) and absence of cellular contamination (calnexin negative). All procedures adhered to MISEV2018¹⁷ and MISEV2023¹¹ guidelines.

### Endothelial Cell Functional Assays

Primary human coronary artery endothelial cells (HCAECs) were treated for 48 hours with EVs isolated from MHL (n=14), MHO (n=9), or MAO (n=16) donors at a standardized dose of 40 µg exosomal protein per 10⁵ cells. Endothelial nitric oxide synthase (eNOS) phosphorylation, ICAM-1 expression, and mitochondrial fusion proteins (MFN1, MFN2) were assessed by immunoblotting. Intracellular reactive oxygen species (ROS) and nitric oxide (NO) were quantified by flow cytometry using DHE and DAF-FM probes, respectively. Mitochondrial ultrastructure was evaluated by TEM.

### Molecular Profiling

EV miRNA cargo was profiled by small RNA sequencing on the Illumina NextSeq 500. Endothelial cell transcriptomic and miRNA responses were assessed by Agilent microarray. Whole blood mRNA and miRNA profiling were performed by RNA sequencing. Plasma proteomic profiling used the Olink Proteomics Explore 3072 platform. Detailed bioinformatics and statistical methods are described in the Online Data Supplement.

### Statistical Analysis

For in vitro experiments, data are presented as mean±SEM. Each data point represents an experiment using cells from a distinct endothelial cell donor exposed to EVs from a different participant. Statistical comparisons were performed using nonparametric unpaired Mann-Whitney U tests. For omics analyses, differential expression was assessed using edgeR (RNA-seq) or limma (microarray) with false discovery rate (FDR) adjustment. Significance was defined as FDR-adjusted P≤0.05. All analyses were conducted using GraphPad Prism (version 10.4.1) and R.

## RESULTS

### Baseline Clinical and Metabolic Profiles

Clinical and metabolic profiles are summarized in Tables 1 and 2. MHO and MAO groups had comparable adiposity, whereas MHL participants were lean. Age and sex distribution did not differ significantly across groups. MAO participants exhibited marked insulin resistance, with HOMA-IR values substantially elevated compared with both MHL and MHO. Blood pressure was significantly elevated in MAO, and 63% of MAO subjects met criteria for hypertension versus none in MHO or MHL. The triglyceride-to-HDL ratio was more than doubled in MAO relative to MHO and MHL. CRP was significantly higher in MAO compared with both MHO and MHL. By contrast, MHO individuals maintained HOMA-IR, blood pressure, lipid, and inflammatory profiles that were clinically comparable to MHL despite equivalent adiposity to MAO. These observations demonstrate that MAO is characterized by a distinct cluster of metabolic abnormalities, whereas MHO retains metabolic features indistinguishable from those of lean individuals.

**Table 2.** Baseline Demographic, Clinical, and Biochemical Characteristics.

| Characteristic | MHL (n=14) |  | MHO (n=9) |  | MAO (n=16) |  | P-Value | P-Value | P-Value |
| --- | --- | --- | --- | --- | --- | --- | --- | --- | --- |
|  | Mean/n | SD/% | Mean/n | SD/% | Mean/n | SD/% | MHL vs MHO | MHL vs MAO | MHO vs MAO |
| Age, y | 43 | 7 | 41 | 6 | 43 | 6 | 0.69 | 0.96 | 0.61 |
| Female, n (%) | 2 | 14% | 4 | 44% | 10 | 63% | 0.26 | 0.02 | 0.65 |
| Male, n (%) | 12 | 86% | 5 | 56% | 6 | 38% |  |  |  |
| BMI, kg/m<sup>2</sup> | 22.4 | 1.2 | 34.7 | 5.9 | 36.9 | 5.2 | <0.001 | <0.001 | 0.37 |
| Waist-to-Hip Ratio | 0.85 | 0.05 | 0.91 | 0.07 | 0.91 | 0.08 | 0.07 | 0.03 | 0.94 |
| Hypertension, n (%) | 0 | 0% | 0 | 0% | 10 | 63% | – | 0.001 | 0.008 |
| SBP, mmHg | 112 | 7 | 115 | 4 | 118 | 13 | 0.18 | 0.09 | 0.36 |
| DBP, mmHg | 68 | 7 | 73 | 7 | 78 | 8 | 0.12 | 0.002 | 0.15 |
| Glucose, mg/dL | 83 | 6 | 86 | 7 | 92 | 8 | 0.35 | 0.002 | 0.06 |
| CRP, mg/dL | 0.01 | 0.01 | 0.02 | 0.01 | 0.04 | 0.02 | 0.06 | <0.001 | 0.009 |
| HOMA-IR | 0.97 | 0.58 | 1.93 | 1.41 | 4.52 | 3.21 | 0.08 | <0.001 | 0.01 |
| TG/HDL | 1.08 | 0.29 | 1.00 | 0.26 | 2.43 | 1.27 | 0.46 | <0.001 | <0.001 |
| HDL, mg/dL | 62 | 8 | 60 | 10 | 53 | 17 | 0.76 | 0.07 | 0.17 |
| LDL, mg/dL | 107 | 36 | 93 | 24 | 124 | 35 | 0.27 | 0.19 | 0.01 |
| TG, mg/dL | 66 | 14 | 60 | 18 | 117 | 47 | 0.44 | <0.001 | <0.001 |
| Education ≤HS, n (%) | 9 | 64% | 7 | 78% | 5 | 31% | 0.82 | 0.15 | 0.07 |
| Current Smoker, n (%) | 9 | 64% | 5 | 56% | 6 | 38% | 0.84 | 0.18 | 0.65 |
| Non-Smoker, n (%) | 4 | 29% | 4 | 44% | 10 | 63% |  |  |  |
| Smoking NR, n (%) | 1 | 7% | 0 | 0% | 0 | 0% |  |  |  |
| Alcohol Use, n (%) | 3 | 21% | 4 | 44% | 4 | 25% | 0.48 | 1.00 | 0.65 |

Continuous variables presented as mean±SD; categorical variables as n (%). P-values by Mann-Whitney U test for continuous variables and Fisher exact test for categorical variables. BMI indicates body mass index; CRP, C-reactive protein; DBP, diastolic blood pressure; HDL, high- density lipoprotein; HOMA-IR, homeostatic model assessment of insulin resistance; HS, high school; LDL, low-density lipoprotein; MAO, metabolically abnormal obese; MHL, metabolically healthy lean; MHO, metabolically healthy obese; NR, not reported; SBP, systolic blood pressure; TG, triglycerides.

### Circulating Markers of Endothelial Activation and Inflammation

Plasma ICAM-1 was significantly elevated in MAO compared with MHL, with intermediate levels observed in MHO (Figure 1A). Circulating CXCL8 (IL-8) was highest in MAO, with modest elevation in MHO relative to MHL (Figure 1C). Leptin concentrations increased progressively from MHL to MHO and MAO, reflecting increasing adiposity-associated metabolic stress (Figure 1B). These findings indicate that MAO is associated with overt endothelial activation and systemic inflammation, while MHO is characterized by low-grade activation that is already detectable at the systemic level.

**Figure 1.**
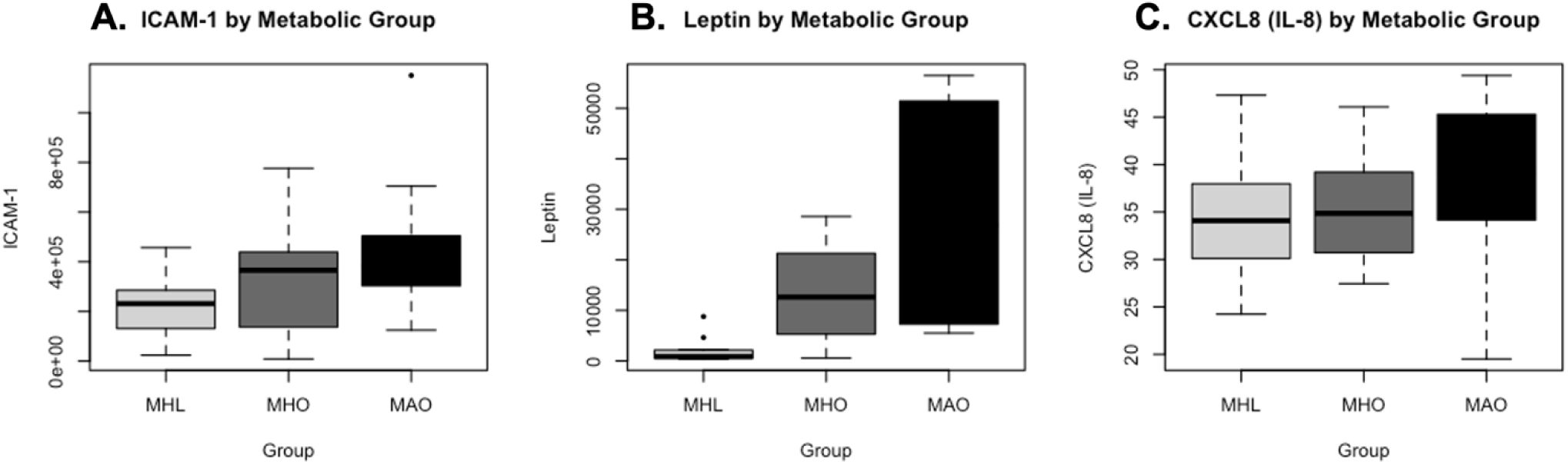
MAO Is Associated With Overt Endothelial Activation and Systemic Inflammation, With Low-Grade Activation Already Detectable in MHO. Box-and-whisker plots showing circulating levels of endothelial and inflammatory markers in metabolically healthy lean (MHL), metabolically healthy obese (MHO), and metabolically abnormal obese (MAO) individuals. (A) Circulating ICAM-1 levels demonstrating a stepwise increase across metabolic phenotypes. (B) Plasma leptin concentrations, markedly elevated in obesity and highest in MAO. (C) Circulating CXCL8 (IL-8) levels showing higher values in MAO compared with MHL and MHO. Boxes represent the interquartile range with the median indicated by the horizontal line; whiskers denote the minimum and maximum values. Statistical comparisons across groups were performed using nonparametric methods. ICAM-1 indicates intercellular adhesion molecule 1; CXCL8, C-X-C motif chemokine ligand 8.

### Plasma-Derived EVs Distinguish Metabolic Phenotypes

TEM revealed round, bilayer vesicles ranging from 50 to 150 nm with characteristic cup-shaped morphology (Figure 2A-C). Immunoblotting confirmed vesicular identity through expression of canonical markers (CD9, CD63, CD81, ALIX) and absence of cellular contamination (Figure 2D). NTA confirmed a unimodal size distribution with mean diameters within the expected small EV range across all groups (Figure 2E-F). Overall particle concentration was elevated in both obese groups relative to lean controls, consistent with increased vesicle secretion linked to adipose and hepatic stress.

**Figure 2.**
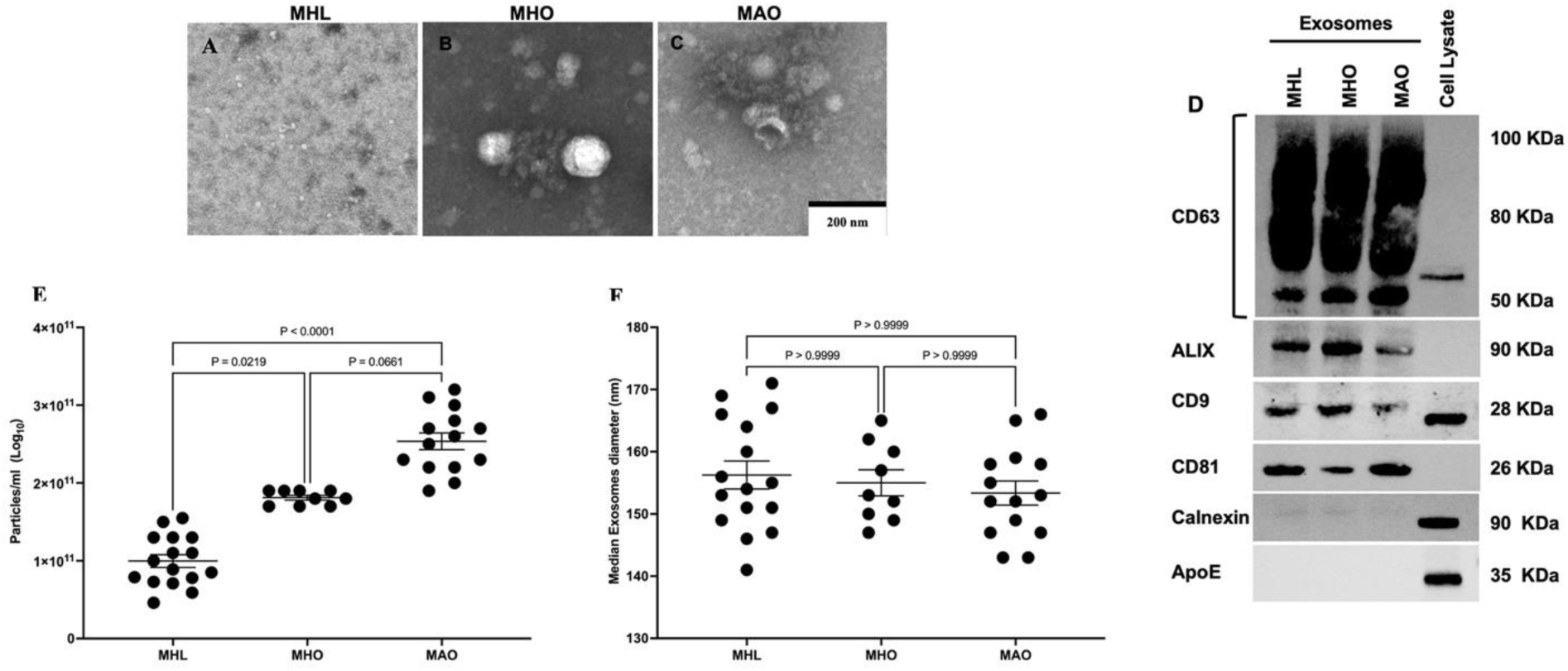
Plasma-Derived Small Extracellular Vesicles Are Successfully Isolated Across All Metabolic Phenotypes. (A-C) Representative transmission electron microscopy (TEM) images showing rounded, bilayer vesicles (50-150 nm) with typical cup-shaped morphology. (D) Immunoblot validation confirming enrichment of canonical exosomal markers CD9, CD63, CD81, and ALIX, with the absence of the endoplasmic reticulum protein Calnexin and ApoE, confirming sample purity (n=3). (E-F) Nanoparticle tracking analysis (NTA) demonstrating comparable mean diameters across groups (MHL ≈141 nm; MHO ≈146 nm; MAO ≈171 nm), all within the expected small-EV size range, with elevated particle concentration in obese groups. Data are expressed as mean±SEM; n=14 (MHL), 9 (MHO), 16 (MAO). Statistical comparisons by Kruskal-Wallis with Dunn’s post-hoc test.

### EV MicroRNA Cargo Defines a Protective-Versus-Pathogenic Signature

Unbiased small RNA sequencing identified 48 differentially expressed miRNAs distinguishing MHO from MAO (FDR-adjusted P≤0.05; Figure 3). MAO-derived EVs were enriched in miR-3613-3p, miR-6842-3p, and miR-326, miRNAs previously implicated in inflammatory signaling and insulin resistance. Pathway enrichment analysis revealed that MAO-enriched miRNA targets mapped to insulin resistance, PI3K-Akt signaling, AMPK suppression, and TNF and Toll-like receptor cascades (Figure 3B).

**Figure 3.**
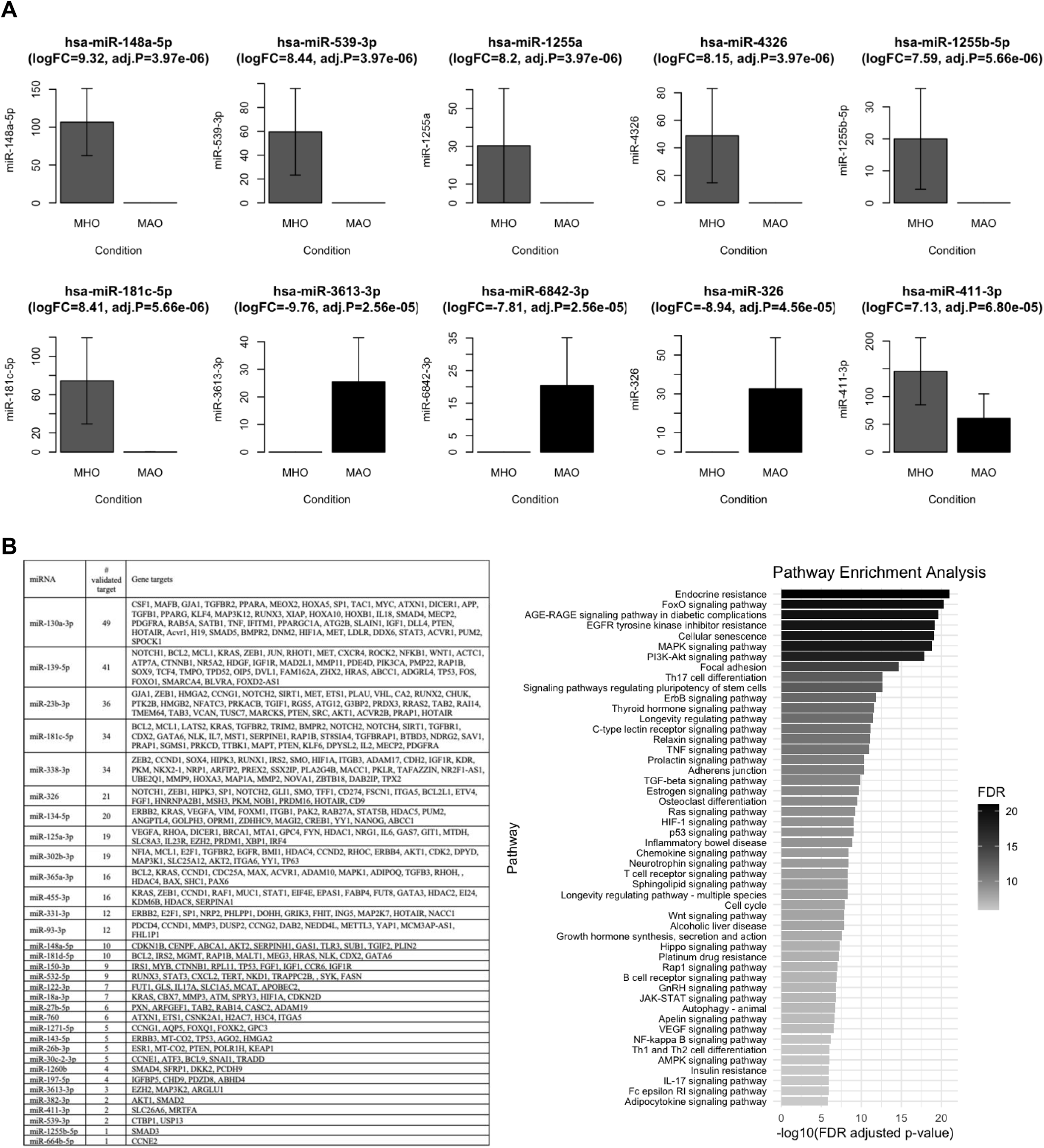
EV MicroRNA Cargo Distinguishes MHO From MAO, With MAO-Enriched MiRNAs Targeting Insulin Resistance, Inflammation, and Oxidative Stress Pathways. (A) Bar plots showing normalized counts of selected plasma-derived EV miRNAs differentially expressed between MHO (grey bars) and MAO (black bars). MicroRNAs displayed were selected based on FDR-adjusted significance and magnitude of differential expression. Log₂ fold-change and adjusted P values (Benjamini-Hochberg correction) are indicated. Bars represent mean±SEM. Statistical analysis by negative binomial modeling with edgeR. (B) KEGG pathway enrichment analysis of validated mRNA targets corresponding to differentially expressed EV miRNAs. Pathways ranked by -log₁₀ FDR-adjusted P value. Enriched pathways include insulin resistance, AGE-RAGE signaling, FOXO signaling, PI3K-Akt signaling, MAPK signaling, inflammatory and immune signaling pathways, and processes related to oxidative stress and cellular senescence. Target interactions were curated from TarBase, miRTarBase, and miRecords. Pathway overrepresentation was evaluated using hypergeometric testing with Benjamini-Hochberg FDR correction. MiRNAs not reaching FDR-adjusted significance (P>0.05) are included for biological context.

In contrast, MHO-derived EVs were enriched in miR-148a-5p, miR-181c-5p, miR-1255a, miR-4326, and miR-539-3p. These miRNAs targeted genes involved in extracellular matrix remodeling, angiogenesis, and TGF-β modulation, consistent with adaptive adipose expansion and vascular protection. Of note, these opposing EV miRNA patterns closely mirror whole-plasma miRNA differences previously reported between MHO and MAO by Rovira-Llopis et al.,20 demonstrating that the circulating miRNA divergence is at least partly vesicle-borne. Together, these findings establish a protective-versus-pathogenic pattern in EV miRNA cargo that distinguishes MHO from MAO.

### MAO-Derived EVs Drive Endothelial Dysfunction; MHO EVs Initiate Early Impairment

To determine whether EV cargo differences translate into functional consequences, HCAECs were treated with plasma-derived EVs. MAO-derived EVs markedly suppressed eNOS phosphorylation (Ser1177) and robustly upregulated ICAM-1 expression compared with both MHL and MHO (Figure 4). Total eNOS protein levels did not differ among groups, indicating post-translational inhibition rather than altered protein expression. MHO-derived EVs induced an intermediate phenotype, with modest but significant reductions in eNOS phosphorylation and increases in ICAM-1 relative to control cells. MHL EVs preserved endothelial homeostasis. These findings delineate a graded continuum of endothelial dysfunction in which MAO EVs drive overt inflammatory activation and MHO EVs initiate early functional impairment.

**Figure 4.**
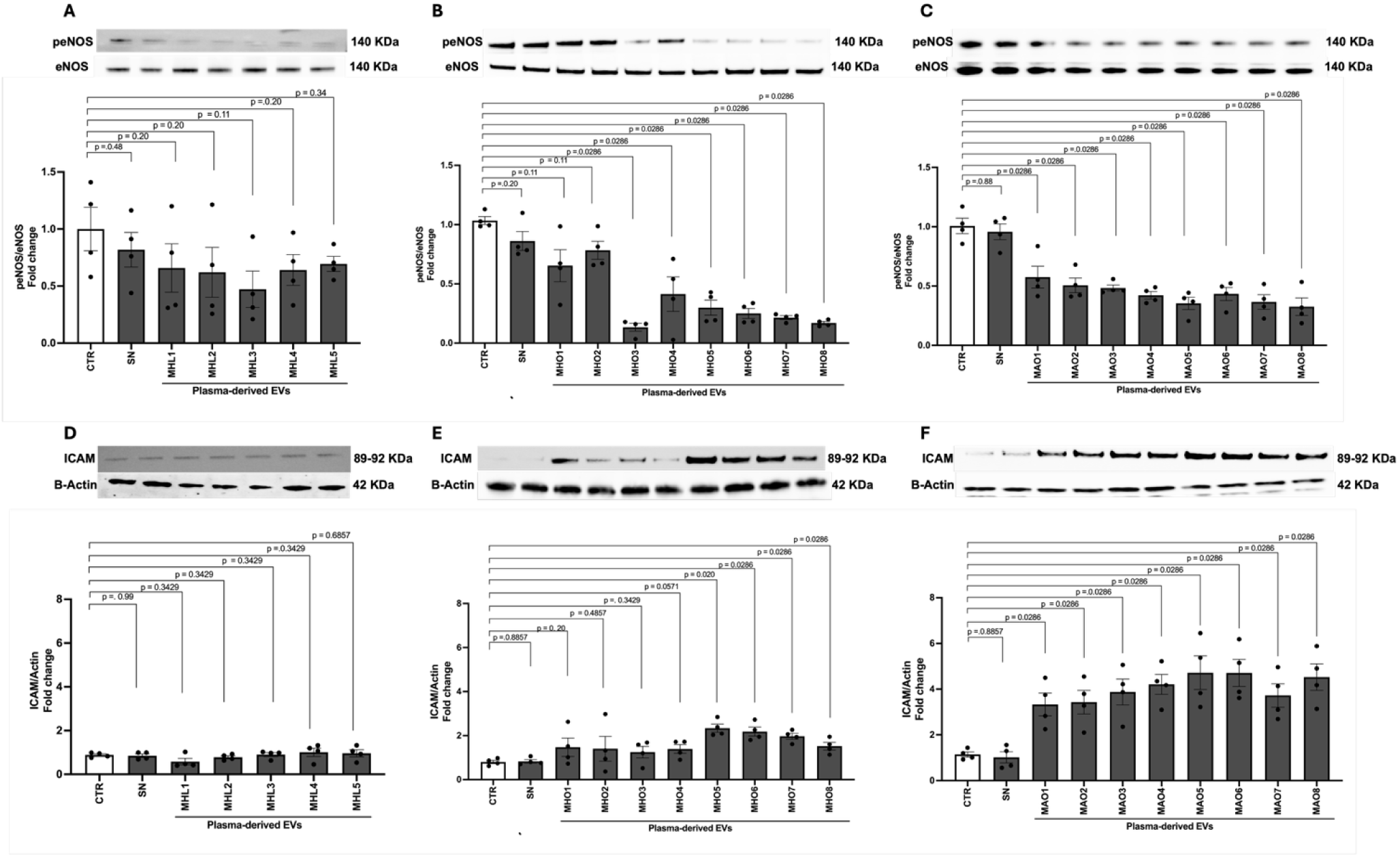
MAO-Derived EVs Drive Overt Endothelial Dysfunction; MHO EVs Initiate Early Impairment. (A-C) Representative immunoblots and quantification of phosphorylated endothelial nitric oxide synthase (p-eNOS, Ser1177) normalized to total eNOS in human coronary artery endothelial cells (HCAECs) following 48-hour exposure to plasma-derived EVs isolated from MHL, MHO, and MAO donors. Untreated control (CTR) and serum-free (SN) conditions are shown for reference. (D-F) Representative immunoblots and quantification of intercellular adhesion molecule-1 (ICAM-1) expression normalized to β-actin under the same conditions. MAO-derived EVs produced marked suppression of p-eNOS and robust induction of ICAM-1, whereas MHO EVs induced intermediate changes consistent with early endothelial dysfunction. MHL EVs had minimal effects. Bars represent mean±SEM from n=4 independent HCAEC donors, with each dot indicating an individual biological replicate. Statistical comparisons by nonparametric Mann-Whitney tests, with exact P values indicated.

### MAO EVs Amplify Endothelial Oxidative Stress and Deplete Nitric Oxide

MAO-derived EVs caused a robust increase in endothelial ROS generation and a pronounced loss of NO bioavailability relative to both MHL and MHO (Figure 5). MHO EVs induced a modest increase in ROS and a moderate reduction in NO relative to control, suggesting early oxidative stress and partial eNOS uncoupling. MHL EVs maintained physiological ROS and NO levels. This pattern mirrors the EV miRNA profiles, where enrichment of oxidative stress-related miRNAs and loss of endothelial-protective miR-126 suggest a mechanistic link between EV signaling and the redox imbalance observed in endothelial cells.

**Figure 5.**
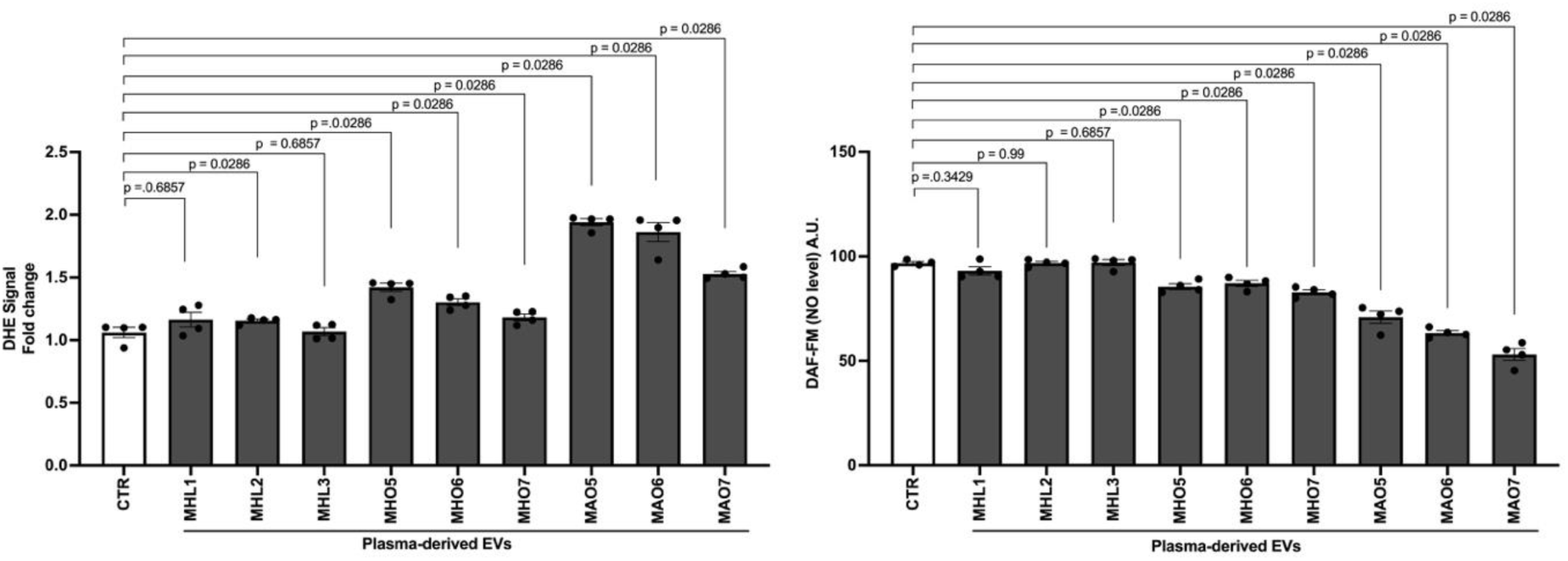
MAO EVs Amplify Endothelial Oxidative Stress and Deplete Nitric Oxide Bioavailability. Bar plots showing intracellular reactive oxygen species (ROS) generation and nitric oxide (NO) bioavailability in HCAECs following 48-hour exposure to plasma-derived EVs from MHL, MHO, and MAO donors. Untreated control (CTR) cells shown for reference. Left panel: Quantification of endothelial ROS production assessed by dihydroethidium (DHE) fluorescence, expressed as fold change relative to control. Right panel: Quantification of endothelial NO levels measured by DAF-FM fluorescence. MAO EVs caused robust ROS elevation and pronounced NO depletion; MHO EVs induced intermediate changes. MHL EVs maintained physiological levels. Bars represent mean±SEM from n=4 independent HCAEC donors. Statistical comparisons by Mann-Whitney tests, with exact P values indicated.

### MAO EVs Disrupt Mitochondrial Integrity; MHO EVs Induce an Intermediate Phenotype

TEM revealed striking ultrastructural differences in endothelial mitochondria following EV exposure (Figure 6A-E). MAO EVs produced swollen, spherical mitochondria with fragmented and disorganized cristae, consistent with mitochondrial stress and loss of respiratory efficiency. MHO EVs induced an intermediate phenotype with occasional cristae irregularities and early swelling. MHL EVs maintained elongated mitochondria with well-defined cristae, indicative of preserved fusion-fission equilibrium. Immunoblot analysis showed a directional decline in mitofusin 1 (MFN1) and mitofusin 2 (MFN2) expression from MHL through MHO to MAO, though pairwise comparisons did not reach statistical significance with the current sample size (n=3 HCAEC donors; Figure 6F-G). Together, these findings demonstrate that EV-mediated suppression of eNOS signaling and redox imbalance converge on mitochondrial dysfunction across the metabolic obesity continuum.

**Figure 6.**
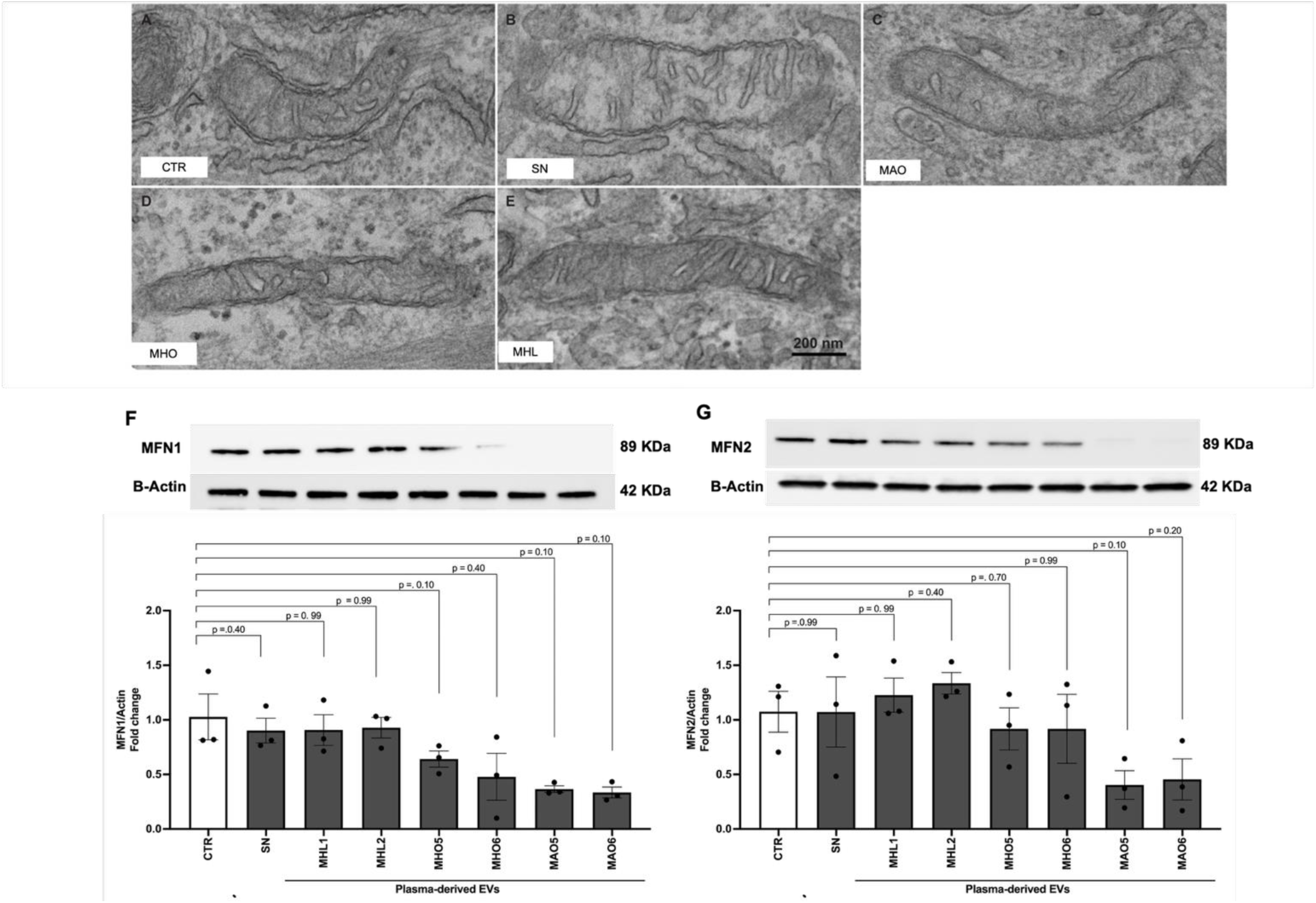
MAO EVs Disrupt Endothelial Mitochondrial Integrity and Suppress Fusion Signaling. (A-E) Representative transmission electron microscopy (TEM) images of HCAECs following 48-hour exposure to plasma-derived EVs. Control- and MHL-EVs treated cells exhibit elongated mitochondria with preserved double membranes and densely organized cristae (A, B, E). MAO EVs exposure induces swollen, rounded mitochondria with disrupted or sparse cristae (C). MHO EVs produces an intermediate phenotype characterized by partially preserved but irregular cristae architecture (D). Scale bar=200 nm. (F-G) Representative immunoblots and densitometric quantification of mitochondrial fusion proteins mitofusin-1 (MFN1) and mitofusin-2 (MFN2) normalized to β-actin. MFN1 and MFN2 expression was preserved in MHL-treated cells, showed modest attenuation following MHO EVs exposure, and was further reduced in response to MAO-derived EVs. Bars represent mean±SEM from n=3 independent HCAEC donors. Statistical comparisons by Mann-Whitney tests.

### MAO-Derived EVs Reprogram the Endothelial MicroRNA and Transcriptional Landscape

MAO-derived EVs induced 50 endothelial miRNAs with robust differential expression (FDR-adjusted P≤0.05 and |log₂FC| > 2), with 36 upregulated and 14 downregulated (Figure 7). Notably altered miRNAs, including miR-1269b, miR-138-2-3p, miR-500a-3p, and miR-4713-5p, have been previously implicated in endothelial activation and inflammatory signaling. The complete list is provided in Supplemental Table S2. By contrast, MHO-derived EVs induced only 2 significantly altered miRNAs (miR-3194-5p upregulated; miR-664a-5p downregulated; Supplemental Figure S4), which did not overlap with the MAO-associated set, indicating that MHO EVs engage distinct and more restricted regulatory pathways.

**Figure 7.**
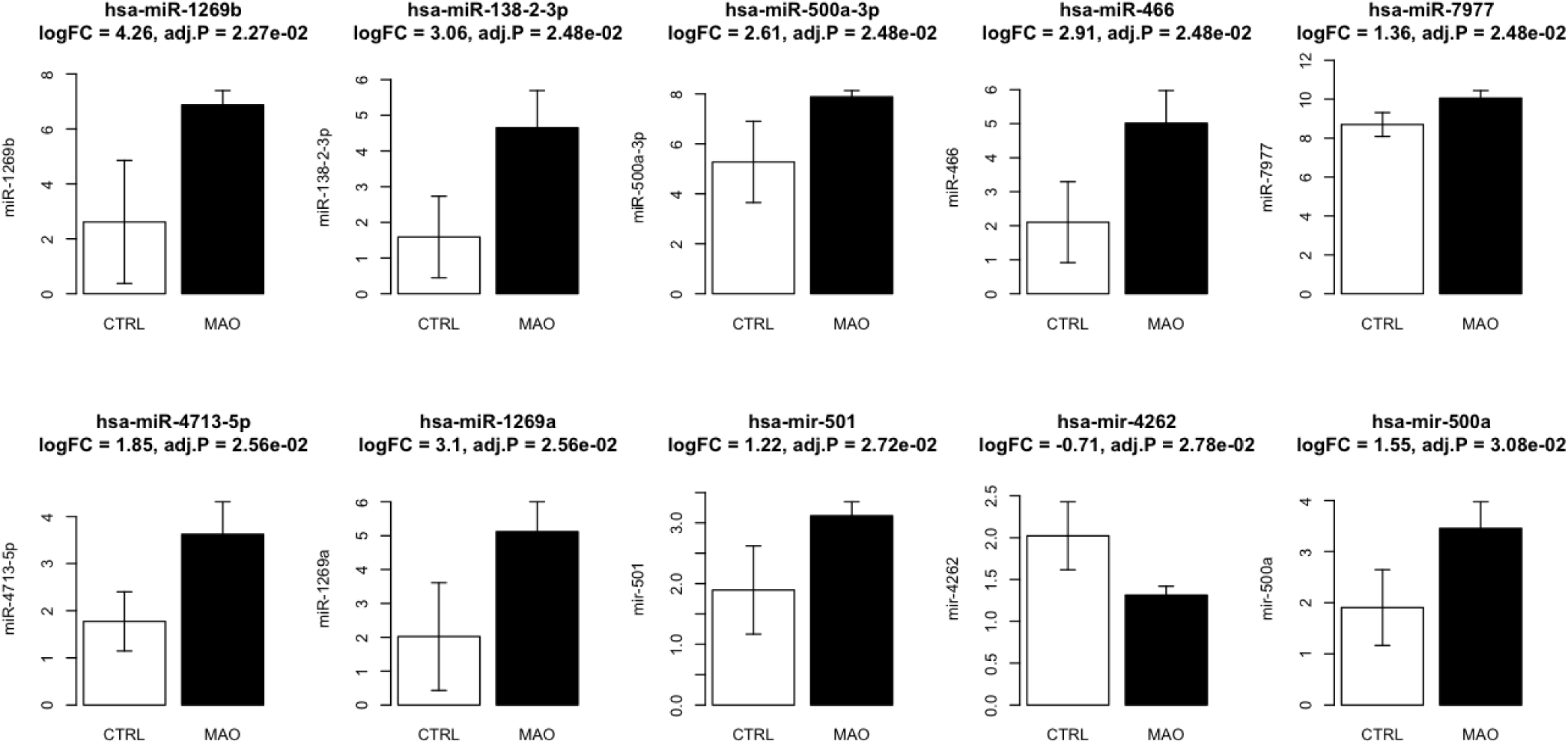
MAO-Derived EVs Induce Broad Endothelial MicroRNA Reprogramming Consistent With Inflammatory Activation. HCAECs were treated for 48 hours with plasma-derived EVs isolated from metabolically abnormal obese (MAO) donors (n=8) or left untreated (CTRL), followed by endothelial microRNA profiling. Shown are selected microRNAs that were significantly differentially expressed in MAO EVs-treated endothelial cells compared with untreated control cells. Bar graphs display mean normalized microRNA expression for each group. Fold change (FC), log fold change (logFC), nominal P values, and FDR-adjusted P values are indicated above each panel. Bars represent data from n=2 independent HCAEC donors. A total of 23 miRNAs were significantly dysregulated (22 upregulated, 1 downregulated); the complete list is provided in Supplemental Table S3.

Consistent with these miRNA changes, endothelial cells exposed to MAO EVs exhibited significantly higher expression of ICAM1 and ICAM3 compared with MHO-treated cells. Expanded analysis of additional adhesion molecules (ICAM4, ICAM5) demonstrated similar directional trends and is provided in the Supplemental Material. These findings demonstrate that endothelial transcriptional reprogramming by obesity-derived EVs is highly dependent on the metabolic phenotype of the donor.

### Systemic Molecular Signatures Corroborate EV-Mediated Findings

At the systemic level, whole blood miRNA sequencing revealed graded molecular alterations across the MHL-MHO-MAO continuum (Supplemental Figure S2). MHO individuals showed modulation of endothelial-regulatory miRNAs, including miR-126-3p, miR-126-5p, and miR-181c-5p, relative to MHL. MAO showed more pronounced dysregulation, with significant downregulation of miR-30a-5p and alterations in miR-192-3p and miR-29a-3p. Whole blood transcriptomic analysis of ICAM1, MFN1, and MFN2 showed minimal variation across groups (Supplemental Figure S3), indicating that whole blood captures only low-amplitude systemic signals. Compared with plasma-derived EV miRNAs, whole blood changes were directionally concordant but of lower magnitude, consistent with the concept that EVs represent a more proximal and enriched signaling compartment of metabolic stress.

## DISCUSSION

The goal of this study was to determine whether circulating EVs mediate graded endothelial dysfunction across the metabolic obesity continuum in adults of African ancestry. The major findings are: (1) MAO-derived EVs carry a pathogenic miRNA signature and drive overt endothelial dysfunction, including suppression of eNOS phosphorylation, depletion of NO, robust oxidative stress, ICAM-1 upregulation, and mitochondrial fragmentation; (2) MHO-derived EVs carry a distinct miRNA profile and already induce measurable impairments in endothelial NO signaling, inflammatory activation, and mitochondrial morphology, albeit to a lesser extent than MAO EVs; and (3) unbiased transcriptomic and proteomic profiling reveal that the molecular divergence between MHO and MAO is encoded within circulating vesicles before overt cardiometabolic disease manifests. Combined, these results demonstrate that MHO is not a vascularly neutral state but rather an active, intermediate stage of EV-mediated endothelial stress (Central Illustration Figure). To our knowledge, this is the first study to directly compare the vascular effects of EVs derived from MHO versus MAO individuals using integrated EV profiling, endothelial functional assays, organelle-level imaging, and transcriptomic analyses.

### MHO Is Not Benign

Clinically, MHO has been characterized by preserved insulin sensitivity and favorable lipid profiles despite excess adiposity. However, longitudinal studies consistently demonstrate progression to MAO with increased cardiovascular risk.22,23 In our study, EVs derived from MHO individuals exhibited a molecular and functional profile distinct from lean controls. In endothelial cells, MHO EVs induced modest but reproducible reductions in eNOS phosphorylation and NO bioavailability, increased ICAM-1 expression, elevated ROS, and partial disruption of mitochondrial morphology. Although these effects were less severe than those observed with MAO EVs, they indicate that vascular homeostasis is already compromised in MHO. MHO should therefore be understood as a dynamic and high-risk state rather than a truly protective phenotype.

### EV Cargo and Mechanistic Insights Into the MHO-MAO Transition

MHO-derived EVs were enriched in miR-181c-5p, miR-539-3p, and miR-148a-5p, each implicated in mitochondrial regulation, endothelial activation, and redox balance. miR-181c-5p has been shown to translocate into mitochondria and impairs complex IV activity, elevating ROS and reducing ATP production in cardiomyocytes and endothelial cells.24,25 miR-539-3p regulates angiogenesis and adhesion pathways and upregulates ICAM-1,26,27 consistent with our observed ICAM-1 induction in MHO-treated HCAECs. The relative depletion of miR-126, abundant in lean controls but reduced in MHO and further diminished in MAO, is noteworthy, as this endothelial-protective miRNA maintains PI3K/Akt-eNOS signaling and vascular repair.28–31 MAO-derived EVs carried a more pathogenic miRNA profile and induced substantially stronger endothelial perturbations. Importantly, EV cargo differences were not merely associative; they translated directly into graded functional and structural endothelial phenotypes in vitro, positioning EVs as active conveyors of metabolic health status.

### EV Effects on Endothelial Dysfunction and Mitochondrial Health

EV-mediated signaling produced a clear hierarchy of endothelial phenotypes. MHL EVs preserved homeostasis with intact eNOS phosphorylation, maintained NO bioavailability, and preserved mitochondrial networks. MAO EVs suppressed eNOS phosphorylation, depleted NO, increased oxidative stress, upregulated ICAM-1, and disrupted mitochondrial morphology with reduced MFN1 and MFN2 expression. No previous in vitro study has directly contrasted the vascular effects of EVs from MHO versus MAO individuals. Prior work has shown that EVs from obese or diabetic mice induce insulin resistance and vascular activation when transferred to lean recipients,32 and that adipose-derived vesicles from unhealthy obesity carry pathogenic miRNAs that impair insulin signaling.14 Our findings extend these observations to humans and reveal that MHO EVs, while less pathogenic, already disrupt eNOS phosphorylation and elevate ICAM-1 expression. These data support a 2-step model: in healthy obesity, EV cargo initiates subtle redox and adhesion changes; as metabolic health declines, pathogenic EV signals dominate, locking in overt endothelial dysfunction and mitochondrial injury.

### EV-Induced Transcriptomic Reprogramming of the Endothelium

MAO EVs induced a robust transcriptomic signature of insulin signaling suppression, mitochondrial downregulation, and inflammatory activation. Pathways related to IRS1, PI3K-Akt, and oxidative phosphorylation were downregulated, while NF-κB-driven inflammatory cascades and adhesion molecule expression were enriched. MHO EVs triggered fewer but notable shifts in genes regulating oxidative stress defenses and extracellular matrix remodeling, consistent with the intermediate functional phenotype observed experimentally. These data confirm that MHO EVs induce molecular perturbations that foreshadow progression to MAO, establishing EV-mediated signaling as a plausible driver of the MHO-MAO transition.

### Systemic Context: Whole Blood Molecular Signatures

At the systemic level, whole blood miRNA and transcriptomic profiling revealed graded molecular alterations across the MHL-MHO-MAO continuum. MHO individuals exhibited intermediate regulatory signatures involving pathways related to inflammation, adhesion, and mitochondrial function. Compared with plasma-derived EV miRNAs, whole blood changes were directionally concordant but of lower magnitude, consistent with the concept that EVs represent a more proximal signaling compartment of metabolic stress. These systemic findings provide in vivo context for the EV-mediated endothelial effects observed experimentally.

## CONCLUSIONS

Collectively, our findings establish extracellular vesicles as active mediators of metabolic health status in obesity. Even in metabolically healthy obesity, EVs carry molecular signals that initiate subtle but measurable endothelial and mitochondrial dysfunction. As metabolic health declines, EV signaling becomes increasingly pathogenic, driving overt vascular injury. This graded EV-mediated communication provides a mechanistic explanation for why MHO is frequently transient and predisposed to progression toward MAO. These data establish EV-associated miRNAs as both biomarkers and mechanistic mediators of endothelial and mitochondrial stress across the metabolic obesity spectrum, providing a foundation for early vascular risk stratification and therapeutic targeting.

## AUTHOR CONTRIBUTIONS

**M. Abbas:** Investigation, methodology, formal analysis, data curation, visualization, writing-review and editing. **C. Bragg:** Investigation, methodology, data curation. **A. Gharib:** Resources, writing-review and editing. **M. Lindsey:** Resources, writing-review and editing. **A.G. Elkahloun:** Resources, methodology, data curation (microarray and sequencing). **A. Gaye:** Conceptualization, supervision, funding acquisition, project administration, writing-review and editing.

## ACKNOWLEDGMENTS

We thank the NHGRI Microarray Core Facility for performing sample processing and hybridization of endothelial cell RNA, the NHLBI Electron Microscopy Core for assistance with transmission electron microscopy imaging, the NIH Intramural Sequencing Center for genomics services and sequencing data, and the NHLBI Flow Cytometry Core for their support in ROS and nitric oxide assays.

## DATA AVAILABILITY

The data that support the findings of this study are available from the corresponding author upon reasonable request.

## FIGURE LEGENDS

**Figure 1. MAO Is Associated With Overt Endothelial Activation and Systemic Inflammation, With Low-Grade Activation Already Detectable in MHO.** Box-and-whisker plots showing circulating levels of endothelial and inflammatory markers in MHL, MHO, and MAO individuals. (A) Circulating ICAM-1 levels. (B) Plasma leptin concentrations. (C) Circulating CXCL8 (IL-8) levels. Boxes represent the interquartile range; whiskers denote the minimum and maximum values. Statistical comparisons were performed using nonparametric methods. ICAM-1 indicates intercellular adhesion molecule 1; CXCL8, C-X-C motif chemokine ligand 8; MHL, metabolically healthy lean; MHO, metabolically healthy obese; MAO, metabolically abnormal obese.

**Figure 2. Plasma-Derived Small Extracellular Vesicles Are Successfully Isolated Across All Metabolic Phenotypes.** (A-C) Representative transmission electron microscopy images showing rounded, bilayer vesicles (50-150 nm). (D) Immunoblotting confirming canonical exosomal markers (CD9, CD63, CD81, ALIX) and absence of calnexin. (E) Nanoparticle tracking analysis demonstrating elevated particle concentration in obese groups. (F) Size distribution confirming expected small EV range across all groups. Data are mean±SEM; n=14 (MHL), n=9 (MHO), n=16 (MAO).

**Figure 3. EV MicroRNA Cargo Distinguishes MHO From MAO, With MAO-Enriched MiRNAs Targeting Insulin Resistance, Inflammation, and Oxidative Stress Pathways.** (A) Bar plots showing normalized counts of selected differentially expressed EV miRNAs between MHO and MAO. Log₂ fold-change and FDR-adjusted P values are indicated. Statistical analysis by edgeR. (B) KEGG pathway enrichment analysis of validated mRNA targets ranked by log₁₀ FDR-adjusted P value.

**Figure 4. MAO-Derived EVs Drive Overt Endothelial Dysfunction; MHO EVs Initiate Early Impairment.** (A-C) Immunoblots and quantification of p-eNOS (Ser1177) normalized to total eNOS. (D-F) Immunoblots and quantification of ICAM-1 normalized to β-actin. Bars represent mean±SEM from n=4 independent HCAEC donors. Statistical comparisons by Mann-Whitney test with exact P values indicated. eNOS indicates endothelial nitric oxide synthase; ICAM-1, intercellular adhesion molecule-1.

**Figure 5. MAO EVs Amplify Endothelial Oxidative Stress and Deplete Nitric Oxide Bioavailability.** Bar plots showing intracellular ROS generation (dihydroethidium fluorescence, left) and NO bioavailability (DAF-FM fluorescence, right) in HCAECs following 48-hour EV exposure. Bars represent mean±SEM from n=4 independent HCAEC donors. Statistical comparisons by Mann-Whitney test. ROS indicates reactive oxygen species; NO, nitric oxide.

**Figure 6. MAO EVs Disrupt Endothelial Mitochondrial Integrity and Suppress Fusion Signaling.** (A-E) Representative TEM images. MAO EVs induce swollen, fragmented mitochondria with disorganized cristae; MHO EVs produce an intermediate phenotype; MHL EVs preserve elongated mitochondria. Scale bar=200 nm. (F-G) Immunoblot quantification of MFN1 and MFN2 normalized to β-actin. Bars represent mean±SEM from n=3 HCAEC donors. Statistical comparisons by Mann-Whitney test. MFN1 indicates mitofusin 1; MFN2, mitofusin 2.

**Figure 7. MAO-Derived EVs Induce Broad Endothelial MicroRNA Reprogramming Consistent With Inflammatory Activation.** HCAECs were treated for 48 hours with MAO-derived EVs (n=8 donors) or left untreated. Selected significantly differentially expressed miRNAs are shown. Fold change, logFC, and FDR-adjusted P values are indicated. Bars represent mean normalized expression from n=2 HCAEC donors. The complete list of differentially expressed miRNAs is provided in Supplemental Table S2.

## Supplemental Figures (in Online Data Supplement)

Supplemental Figure S1. EV miRNA-target networks and validated mRNA target table.

Supplemental Figure S2. Whole blood microRNA profiling across metabolic obesity phenotypes (miR-126-3p, miR-126-5p, miR-181c-5p, miR-30a-5p, miR-192-3p, miR-29a-3p).

Supplemental Figure S3. Systemic transcriptional signatures of ICAM1, MFN1, and MFN2 in whole blood.

Supplemental Figure S4. Endothelial microRNAs differentially expressed following MHO-derived EV exposure (miR-3194-5p, miR-664a-5p).

Supplemental Figure S5. Endothelial adhesion gene expression (ICAM1, ICAM3, ICAM4, ICAM5) following MHO- and MAO-derived EV exposure.

## CENTRAL ILLUSTRATION

**Central Illustration.**
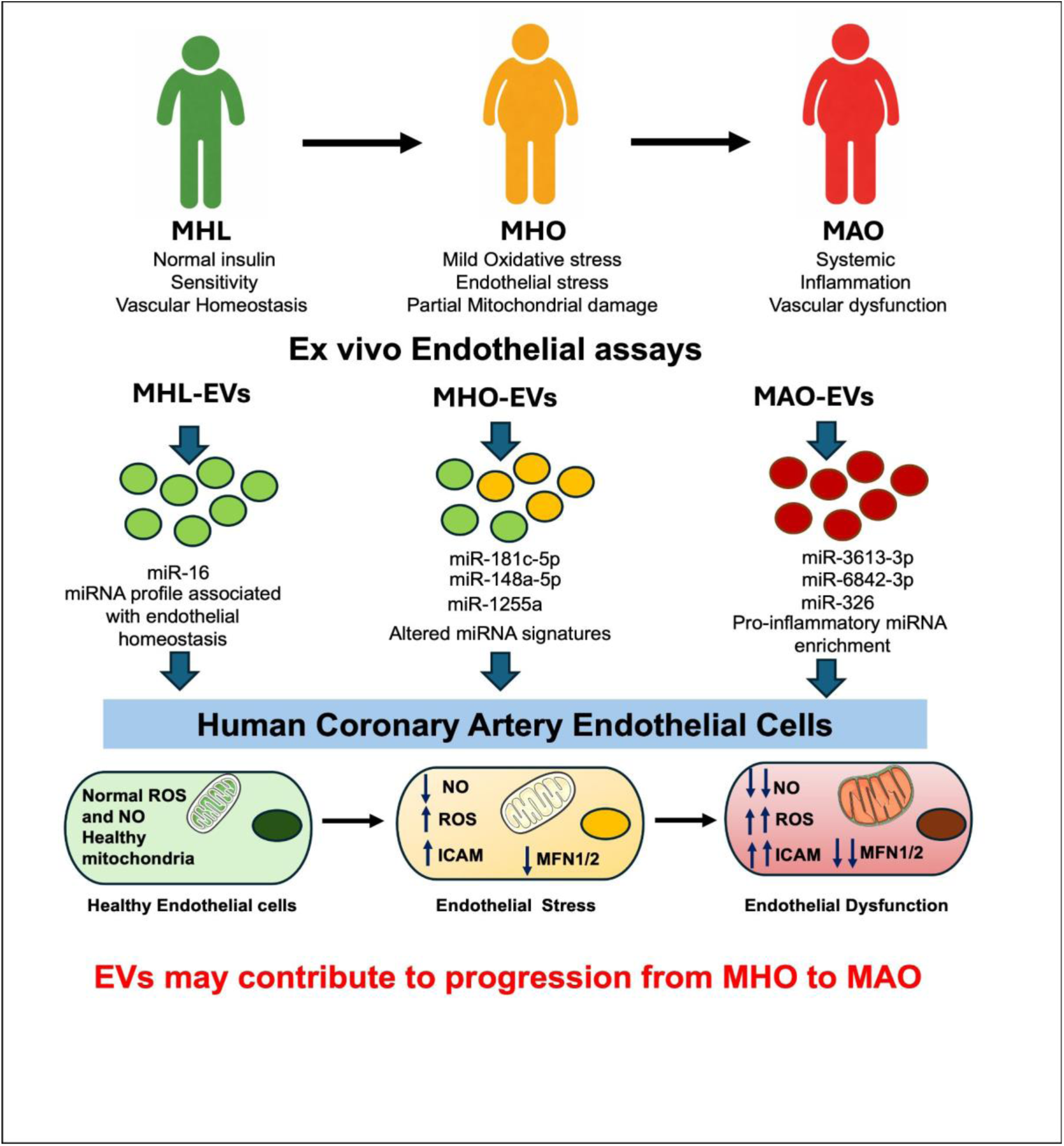
Graded extracellular vesicle-mediated endothelial dysfunction across the metabolic obesity continuum. Circulating extracellular vesicles (EVs) from metabolically healthy lean (MHL), metabolically healthy obese (MHO), and metabolically abnormal obese (MAO) individuals carry distinct microRNA cargo that induces graded endothelial responses in human coronary artery endothelial cells. MHL-derived EVs preserve endothelial homeostasis with normal nitric oxide (NO) bioavailability, low reactive oxygen species (ROS), and intact mitochondria. MHO-derived EVs, enriched in miR-181c-5p, miR-148a-5p, and miR-1255a, initiate early endothelial stress characterized by modest NO reduction, ROS elevation, ICAM-1 upregulation, and partial mitochondrial disruption. MAO-derived EVs, enriched in pro-inflammatory miRNAs including miR-3613-3p, miR-6842-3p, and miR-326, drive overt endothelial dysfunction with pronounced NO depletion, robust oxidative stress, inflammatory activation, and mitochondrial fragmentation with reduced MFN1/2 expression. These findings demonstrate that MHO is not a benign state but an intermediate stage of EV-mediated vascular stress that may predispose to progression toward MAO.

## Nonstandard Abbreviations and Acronyms

MHO: metabolically healthy obesity
MAO: metabolically abnormal obesity
MHL: metabolically healthy lean
EVs: extracellular vesicles
miRNA: microRNA
eNOS: endothelial nitric oxide synthase
NO: nitric oxide
ROS: reactive oxygen species
ICAM-1: intercellular adhesion molecule-1
MFN1/2: mitofusin 1/2
HCAECs: human coronary artery endothelial cells
HOMA-IR: homeostatic model assessment of insulin resistance
CRP: C-reactive protein
TEM: transmission electron microscopy
NTA: nanoparticle tracking analysis

